# Shifts in antimalarial drug policy since 2006 have rapidly selected *P. falciparum* resistance alleles in Angola

**DOI:** 10.1101/2020.09.29.310706

**Authors:** Emily R Ebel, Fátima Reis, Dmitri A Petrov, Sandra Beleza

**Affiliations:** Department of Biology, Stanford University, Stanford, CA 94305, United States of America; Department of Pediatrics – Infectious Disease, Stanford University School of Medicine, Stanford, CA 94305, United States of America; Hospital Regional de Cabinda, C5QW+XP Cabinda, Angola; Department of Genetics, University of Leicester, LE1 7RH, Leicester United Kingdom

**Keywords:** *Plasmodium falciparum*, Angola, chloroquine, lumefantrine, drug resistance, selection

## Abstract

**BACKGROUND:** *Plasmodium falciparum* resistance to chloroquine (CQ), the most widely used antimalarial drug, has historically posed a major threat to malaria control in Angola and throughout the world. Although Angola replaced CQ with artemisinin combination therapy (ACT) as a frontline treatment in 2006, malaria cases and deaths have recently been rising. CQ-resistance mutations may still be a contributing factor, given that (1) some also modulate resistance to ACT partner drugs and (2) ACT is not yet consistently implemented across Angola. It is important to continue monitoring all known resistance alleles in *P. falciparum*, but no studies have done so in Angola since 2012.

**METHODS:** We sampled *P. falciparum* DNA from the blood of 50 hospital patients in Cabinda, Angola in 2018. Each infection was genotyped for 13 alleles in the genes *crt, mdr1, dhps, dhfr*, and *kelch13*, which collectively confer resistance to six common drugs. To analyze frequency trajectories over time, we also collated *P. falciparum* genotype data published from across Angola in the last two decades.

**RESULTS:** The two most important alleles for CQ resistance, *crt* 72-76CVIET and *mdr1* 86Y, have both declined in frequency from respective highs of 98% in 1999 and 73% in 2003. However, the former remains at 71% frequency in this sample while the latter has dropped to just 7%. Of seven possible alleles for sulfadoxine-pyrimethamine (SP) resistance in *dhps* and *dhfr*, the average total number per isolate increased from 2.9 in 2004 to 4.4 in 2018. Finally, we detected no non-synonymous polymorphisms in *kelch13*, which is involved in artemisinin resistance in Southeast Asia.

**CONCLUSIONS:** Changes in drug policy in Angola since 2006 appear to have exerted strong selection on *P. falciparum* drug resistance alleles. Resistance to CQ is declining, but due to functional tradeoffs and novel selection at *mdr1* loci, resistance to ACT partner drugs appears to be rising. More haplotype-based studies at *mdr1* will be needed to understand the changing efficacy of multiple drugs. Finally, SP resistance has jumped rapidly since 2014, consistent with widespread use of intermittent SP treatment during pregnancy. These data can be used to support effective drug policy decisions in Angola.

## BACKGROUND

Antimalarial drugs have long been important tools for malaria control^1^. However, their efficacy is constantly threatened by the evolution of drug resistance in *Plasmodium falciparum*^2^. Multiple *P. falciparum* genes are involved in drug resistance, and selection on them varies by allele, genetic background, and drug environment^3–5^. Therefore, frequent monitoring of resistance alleles is crucial to predicting the spread of drug resistance. This is especially true in the West African country of Angola, where malaria cases and deaths are on the rise^6^.

The first anti-malarial drug to enjoy widespread use in Angola was chloroquine (CQ) in the 1950s^7^. CQ resistance was first confirmed in Angola in the 1980s, and by the early 2000s, CQ failure rates exceeded 80%^8,9^. As a result, CQ was discontinued in Angola in favor of artemisinin-based combination therapy (ACT) starting in 2006^10^. To discourage the evolution of artemisinin resistance, artemisinin is used in combination with the longer-acting partner drugs lumefantrine (LMF) or amodiaquine (AQ), which is chemically related to CQ^11^. Artemisinin resistance has not yet appeared in Angola, although many resistant *kelch13* mutations have emerged in Southeast Asia^5,12^. Nonetheless, occasional ACT treatment failures have been reported in Angola due to partner drug resistance^10^.

Strong *P. falciparum* resistance to CQ and AQ is caused by *crt* K76T, a lysine to threonine substitution at codon 76 of the chloroquine resistance transporter (Table 1). A meta-analysis found this allele to be 7.2-fold overrepresented in CQ treatment failures^13^, reflecting its selection by CQ and AQ in many clinical studies (Table 1). In Angola, K76T is found on the haplotype *crt* 72-76 CVIET, which is of Asian origin^14^. CQ resistance has also evolved independently through the haplotype *crt* 72-76 SVMNT in South America and Papua New Guinea^15^.

**Table 1.**
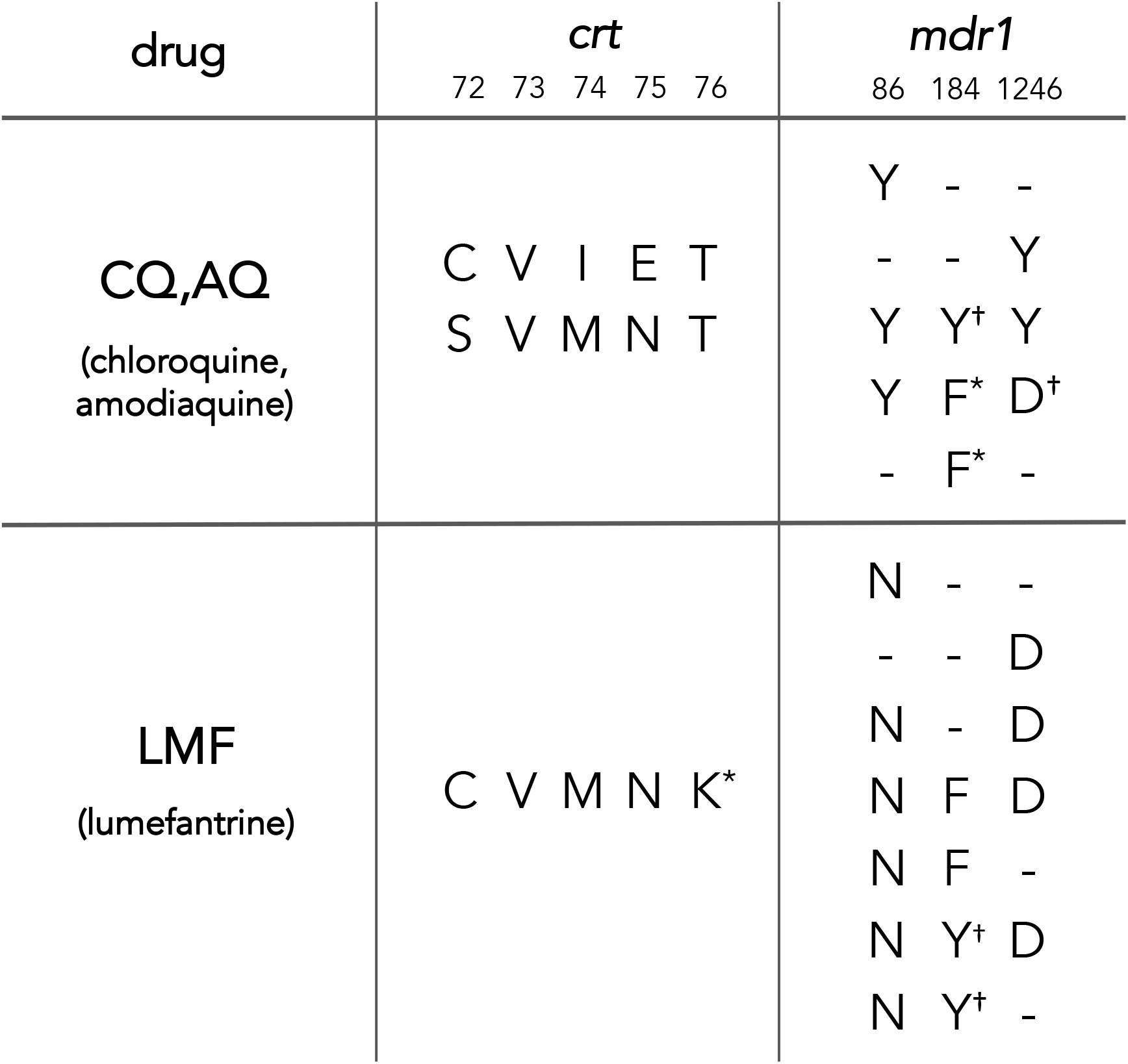
Alleles of *P. falciparum* genes *crt* and *mdr1* that are preferred in the presence of frontline antimalarial drugs. Although CQ has been discontinued in Angola, CQ-resistance loci are also involved in resistance to the ACT partner drugs AQ and LMF. Numbers in the header indicate amino acid position. For *mdr1*, incomplete haplotypes are shown as reported in the literature. *These alleles are unlikely to confer resistance directly, but they are less deleterious than the alternate allele in the presence of drug. *This allele is unlikely to confer resistance directly, but it is linked to other functional alleles. Additional details and references are available in Table S1.

The N86Y allele of *mdr1*, or multidrug resistance gene 1, also confers resistance to CQ and AQ^13^. Although this specific polymorphism dominated early studies of *mdr1* and CQ resistance, the evolution of *mdr1* is complicated by linkage between position 86 and other functional polymorphisms^16^. Precise *mdr1* haplotypes vary among *P. falciparum* populations and drug settings, but in Angola alone, at least six alleles at three *mdr1* positions have been proposed to modulate resistance to CQ, AQ, and the ACT partner drug lumefantrine (LMF) (Table 1; Table S1).

The drug sulfadoxine-pyrimethamine (SP) has also been in widespread use in many African countries since the 1960s^17^. *P. falciparum* quickly began evolving partial resistance to SP, mediated by numerous substitutions in *dhps* and *dhfr*^18^. The risk of SP treatment failure increases with the number of mutant alleles present, with “quintuple mutants” at codons 437/540 of *dhps* and codons 51/59/108 of *dhfr* of particular concern^19–21^. By the early 2000s, these alleles were common in Angola and 25-39% of *P. falciparum* infections failed to respond to SP treatment^9^. SP has since been discontinued as a frontline therapy, but it is still administered to pregnant women to reduce common complications from malaria^22^. Although this approach is generally still useful in Africa^23,24^, its efficacy is waning as additional *dhps* mutations continue to emerge^18,25–27^. In one recent example from Tanzania, a novel mutation at *dhps* 581 was both selected by SP treatment and associated with worse pregnancy outcomes^28^. Because SP is still in widespread use, it is critical to continue monitoring its effectiveness along with variation in its target genes.

In this work, 50 *P. falciparum* infections from Cabinda, Angola were genotyped for 13 markers of drug resistance in the genes *crt*, *mdr1, dhps, dhfr*, and *kelch13*. Similar allele frequency data were also gathered from studies published on Angolan *P. falciparum* in the last two decades. For every locus but *kelch13*, we found temporal patterns of allele frequency change that are consistent with changes in drug policy. This work can inform future decisions on drug administration in Angola, particularly given rapid increases in SP resistance.

## RESULTS

### Genotyping success and MOI

Each sample was successfully genotyped at an average of 12 out of 13 loci (Table S3). The *kelch13* locus had the highest success rate (100%), while *crt* had the lowest success rate (78%). Although the *crt* primers have performed well on other Angolan samples^29^, in this cohort, even the nested protocol amplified products of multiple sizes (Fig S1).

Fifteen of 50 samples had sequence diversity (i.e., peaks of two bases) in at least one resistance marker site. Assuming that double peaks indicated the presence of two strains, the overall multiplicity of infection (MOI) was 1.3.

### Very little polymorphism in *kelch13*

No *kelch13* polymorphisms were observed at codons 578-580, which have been associated with ACT resistance in Southeast Asia and Uganda^12^. Moreover, with the exception of one synonymous variant in one sample, no polymorphism was observed across all 261 *kelch13* codons sequenced in this study.

### Markers of CQ resistance and LMF susceptibility are declining

The CVIET haplotype at *crt* codons 72-76, which confers strong resistance to CQ, was detected at 71% frequency in this study (Fig 1). This represents a significant decline from a peak of 98% in 1999 (*p* = 0.03), although individual estimates have been noisy over time.

**Figure 1.**
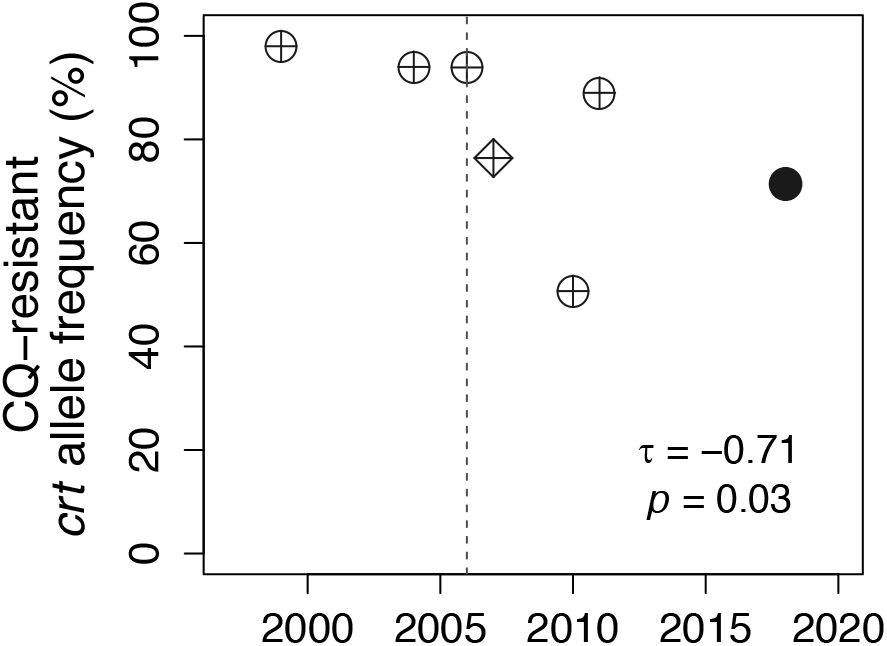
The frequency of CQ (chloroquine)-resistant alleles in *crt* has declined in Angola since the late 1990s. The solid circle indicates new data from this study. Historical data were obtained from^29,31–35^. The diamond indicates the combined presence of two resistant haplotypes (CVIET and SVMNK) at *crt* 72-76 in^29^; all other points represent CVIET only. τ is the Kendall rank correlation between time of sampling and frequency of resistance. The dashed line shows the year that CQ was officially discontinued in Angola.

The *mdr1* allele 86Y, which also confers resistance to CQ, was detected at just 6.5% frequency in this study (Fig 2A). This marker has declined rapidly and steadily from ~80% frequency in 2003 (*p* = 0.007). Accordingly, the alternate allele 86N—which is both CQ-sensitive and LMF-resistant (Table 1) — has increased in frequency to 93.5% (Fig 2B, *p* = 0.006). The linked polymorphism *mdr1* 184F, which is also preferred in the presence of LMF (Table 1), has been rising in frequency at a similar rate (Fig 2B, *p* = 0.087), although it remains less common than 86N. A single sample contained the additional CQ-resistance allele *mdr1* 1246Y (Table 1), which occurred on an 86Y/184Y background (Table S3).

**Figure 2.**
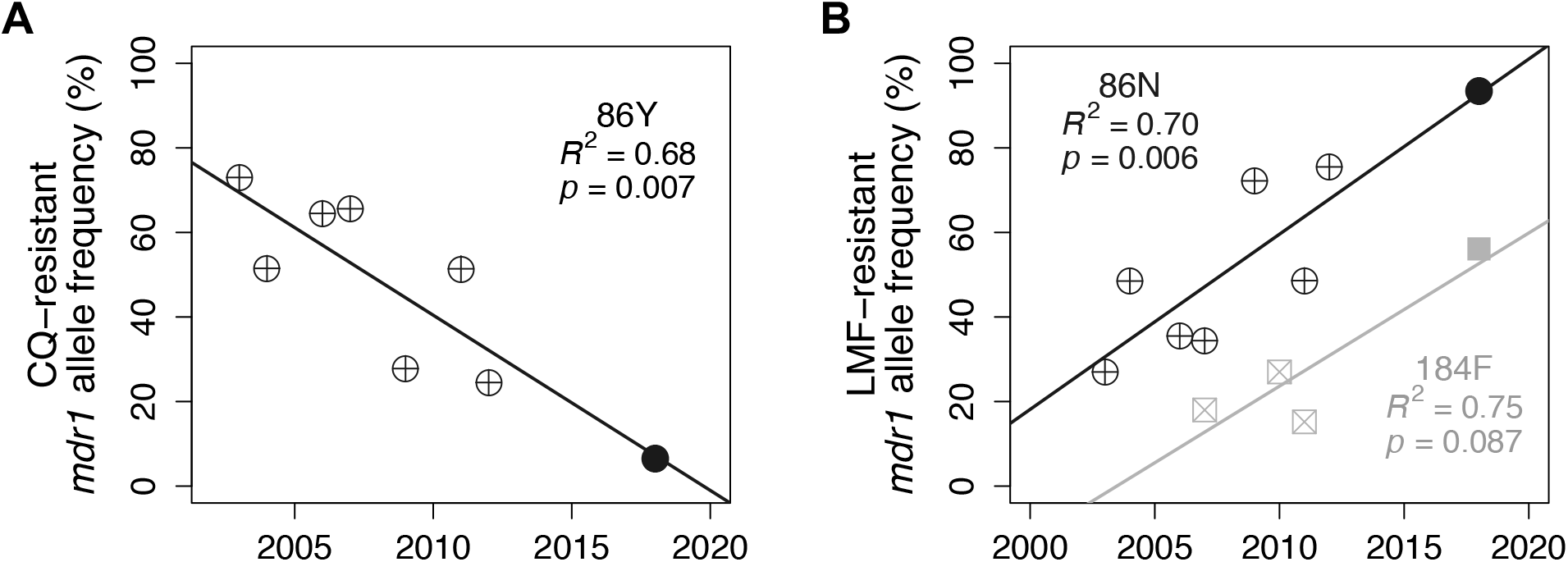
*mdr1* allele frequencies have changed steadily in Angola since the early 2000s. Solid points indicate new data from this study. Historical data were obtained from^29,32–37^. Lines of best fit, variance explained (R^2^), and *p*-values from linear regression are shown for each allele.

### Markers of SP resistance markers have become more common

One third (33%) of *P. falciparum* isolates sampled here were “quintuple mutants” for five *dhfr* and *dhps* alleles that confer strong SP resistance (Fig 3). Compared to samples from migrant workers collected around 2014^38^, this represents a 2.8X increase of quintuple mutants in Angola in less than five years. Three “sextuple mutants” were also observed for the first time in Angola, including resistance alleles at *dhps* codons 436 (22% frequency) and 581 (8.2% frequency). The average number of combined *dhfr/dhps* resistance alleles per isolate has increased sharply over time, from 2.9/7 in 2004 to 4.4/7 in this study (*t* = −9.71, *p* < 2.2 x10^-16^).

**Figure 3.**
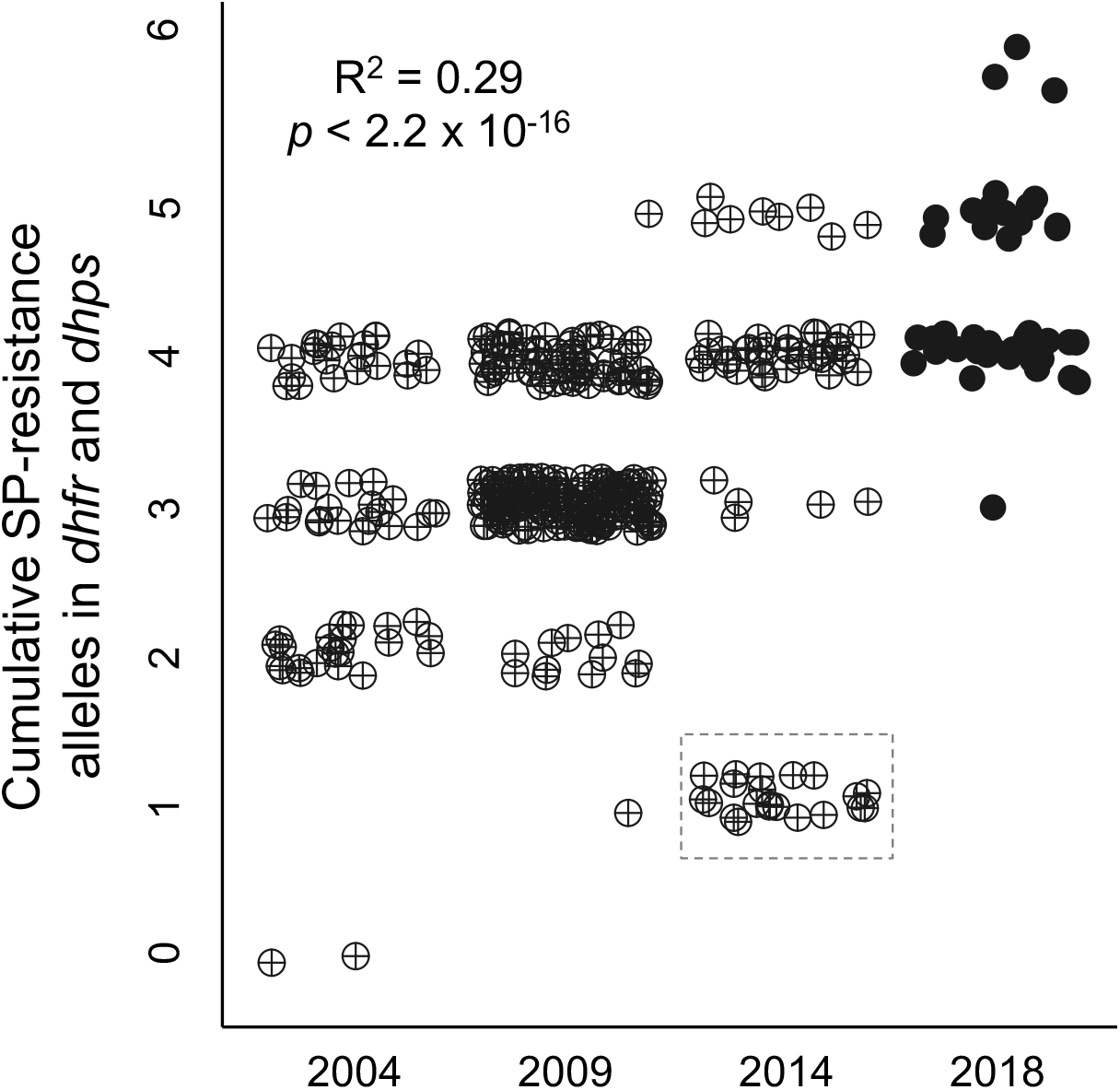
SP-resistant alleles in *dhfr* and *dhps* have become more common in Angola since the early 2000s. Each point represents the total number of resistance allels in both genes from a single isolate. Solid points indicate new data from this study; historical data are from^32,38,41^. Points are jittered horizontally and vertically for clarity. Resistance alleles were counted at *dhfr* codons 51, 59, and 108 and *dhps* codons 436, 437, 540, and 581. The dashed box surrounds a subset of 2014 isolates that carried 0-3 mutant alleles, but for which more precise data were not available. Variance explained (R^2^) and *p*-value are shown are from a linear regression excluding all 2014 data.

## DISCUSSION

The official withdrawal of CQ in Angola since 2006 has likely contributed to the decline of CQ-resistance alleles in *crt* (Fig 1) and *mdr1* (Fig 2A). This result is similar to other African countries that have discontinued CQ, including Malawi, the Gambia, Kenya, Ethiopia, Tanzania, and Grand Comore^39^. In Malawi, clinical CQ sensitivity largely returned after the prevalence of *crt* K76T declined from 85% in 1992 to 13% in 2000^40^. In Angola, however, the prevalence of the CVIET haplotype in Angola remains high at 71% (Fig 1). Although the exact fraction of resistant parasites may vary by locality (Fig 1), these results imply that CQ resistance via *crt* is still standard in Angola. In contrast, the rate of decline of *mdr1* 86Y—the second-most important CQ-resistance allele—is sharp enough to suggest its disappearance from Angola within a few years (Fig 2A). This stark difference between the evolution of *crt* and *mdr1* may be best explained by the differential effects of LMF, an ACT partner drug, on their CQ-sensitive alleles (Table 1). Specifically, wild-type *crt* K76 is only passively selected in the absence of CQ, while wild type *mdr1* N86 directly confers LMF resistance (Table 1). LMF resistance is highest when *mdr1* N86 co-occurs with *mdr1* 184F, which is rapidly spreading in Angola (Fig 2B), and *mdr1* 1246D, which is nearly fixed (Table S3). Consequently, the rise of LMF resistance could soon challenge the success of ACT as currently implemented in Angola.

The most rapid change observed in this study was the increase in total SP-resistance alleles per isolate (Fig 3). Similar increases over time have been reported in a number of other African countries^42–44^, likely in response to the implementation of WHO recommendations for SP during pregnancy. It is now clear that more than five mutations in *dhfr/dhps* contribute to SP resistance: in our sample, seven such mutations were present, and three infections (7.0%) carried six of them. Because intermittent SP treatment currently recommended by the WHO does not eliminate parasitemia^45^, it is strongly expected to select for additional SP resistance. The benefits of SP in pregnancy have outweighed these costs in the past, but the present rate of resistance evolution implies that these benefits may be eroding^27,28^. Further research will be required to weigh the impact of novel resistance haplotypes against other factors impacting SP treatment efficacy. To help accomplish this goal, we emphasize the importance of reporting complete haplotype information for all combined *dhfr/dhps* alleles in each sampled infection.

Finally, we detected no signs of artemisinin-resistant alleles in *kelch13*. This result is consistent with the high efficacy of ACT in Angola^10^, and overall, there is little evidence that artemisinin resistance alleles are spreading in Africa^46^. Monitoring of *kelch13* in Africa is nonetheless important, as artemisinin is the only drug for which resistance alleles are not already widespread.

## CONCLUSIONS

Changes in drug policy since 2006 have had clear impacts on the frequencies of several drug resistance alleles in Angola. Markers of SP resistance are rapidly becoming more common, which endangers the efficacy of intermittent treatment during pregnancy. Resistance to CQ is declining, but resistance to LMF appears to be rising. More frequent monitoring and drug policy adjustments will likely be necessary to regain control of *P. falciparum* malaria in Angola.

## METHODS

### Sample collection and ethics statement

Patients reporting to the Hospital Regional de Cabinda in 2018/2019 with fever, chills, or other malaria symptoms were offered the option to be consented to this study. Sample collection followed protocols approved by Stanford University (IRB #39149) and the Medical Ethics Committee of the University 11^th^ of November in Cabinda. Consented participants’ blood was drawn from a vein and screened under a microscope for *P. falciparum* parasites. If positive, whole blood was filtered through cellulose columns to remove leukocytes^47^. The filtered red blood cells were spotted on Whatman FTA cards (Sigma Aldrich), dried, and stored for at least 6 months.

### DNA extraction and genotyping

To elute DNA, saturated circles were cut out of the Whatman FTA cards and incubated in 800 uL TE buffer (10 mM Tris-Cl, 1 mM EDTA, pH 8.0) with 20 uL Proteinase K (Invitrogen) for 2 hours at 65°C. DNA was extracted from the liquid supernatant using a phenol-chloroform protocol with phase-lock gel tubes^48^.

PCR amplification of the *P. falciparum* genes *crt, mdr1, dhfr, dphs*, and *kelch13* was performed with previously published primers^29,49,50^. Cycling protocols were based on manufacturer recommendations for OneTaq Hot Start 2X Master Mix (NEB) and/or Phusion High-Fidelity PCR Master Mix with HF Buffer (NEB) (Table S2). Reactions were visualized in 1% agarose gels, and if successful, cleaned with ExoSAP-IT (ThermoFisher) and Sanger sequenced (Elim Bio). Sanger chromatogram data were compared to PlasmoDB reference *P. falciparum* sequences using Benchling. Amino acid substitutions were identified in the following positions: *mdr1* 86, 184, and 1246; *crt* 72-76; *dhfr* 50, 51, 59, and 108; *dhps* 436, 437, 540, and 681; and *kelch13* 578-580.

### MOI and allele frequency calculations

For each sample, a double infection was inferred if the sequencing chromatogram showed equally sized, double peaks for any of the 13 analyzed loci. Multiplicity of infection (MOI) was calculated as the total number of infections divided by the total number of samples, as previously described^51^. Similarly, the frequency of each allele was determined based on the total number of infections, with double infections at any locus contributing two genotypes at every locus. Samples without missing data at *dhfr* or *dhps* were also assessed for the presence of up to seven SP-resistance alleles (*dhfr-51I, dhfr-59R, dhfr-108G, dhps-436, dhps-437G, dhps-540E, dhps*-581)^52,53^.

### Collection of historical data

Publications reporting allele frequencies for drug-resistance loci anywhere in Angola since 1995 were gathered from the Worldwide Antimalarial Resistance Network (WWARN) Molecular Surveyor tool (http://www.wwarn.org/molecularsurveyor/), facilitated by a recent review^7^. The original data published in these studies were used to calculated alleles frequencies as described above. For studies that spanned multiple years, the average year was used for time-course analysis (below). Studies that did not provide linked data for *dhfr* and *dhps* (e.g., reported the two genes separately) could not be included.

### Statistical analysis

To evaluate changes in *mdr1* and *dhfr/dhps* alleles over time, linear models were fit to the frequency or count data using the lm function in R. To avoid a bias from incomplete data, all 2014 samples were excluded from the *dhfr/dhps* timecourse analysis. For *crt*, the relationship between CVIET frequency and time was not linear; therefore, Kendall’s rank correlation was applied using the cor.test function in R.

## DECLARATIONS

### Ethics approval and consent to participate

Ethics approval for this study was obtained from Stanford University IRB (#39149) and the Medical Ethics Committee of the University 11^th^ of November in Cabinda.

### Consent for publication

Prior to participation, all study subjects and/or their parents consented in writing to the publication of study results in the scientific literature.

### Author’s Contributions

E.R.E., D.A.P., and S.B. designed the study. F.R. and S.B. supervised the study. E.R.E. collected data, analyzed data, and wrote the manuscript. All authors have approved the final manuscript.

## Acknowledgements

We are immensely grateful to the study participants and staff of the Hospital Regional de Cabinda. Logistic support in sample collection was provided by Dr. Maria das Dores Sungo and Dr. Francisco Casimiro Lubalo, Rector and Vice-Rector of the Faculty of Medicine, University 11th of November, Cabinda. We also thank Rachael Madison, Bàrbara Baro Sastre, and Elizabeth Egan for their assistance in preparing and testing sampling materials.

