## Supplementary material for "Shifts in antimalarial drug policy since 2006 have rapidly selected *P. falciparum* resistance alleles in Angola": SI: Supp Figs and Legends.docx

**Table S1. Evidence and references supporting summary in Table 1.**

1.	Djimdé, A. *et al.* A Molecular Marker for Chloroquine-Resistant Falciparum Malaria. *N. Engl. J. Med.* **344**, 257–263 (2001).

2.	Picot, S. *et al.* A systematic review and meta-analysis of evidence for correlation between molecular markers of parasite resistance and treatment outcome in falciparum malaria. *Malar. J.* **8**, 89 (2009).

3.	Holmgren, G. *et al.* Amodiaquine resistant Plasmodium falciparum malaria in vivo is associated with selection of pfcrt 76T and pfmdr1 86Y. *Infect. Genet. Evol.* **6**, 309–314 (2006).

4.	Ursing, J., Kofoed, P.-E., Rodrigues, A., Rombo, L. & Gil, J. P. *PLASMODIUM FALCIPARUM GENOTYPES ASSOCIATED WITH CHLOROQUINE AND AMODIAQUINE RESISTANCE IN GUINEA-BISSAU*. (2007).

**Figure S1. PCR with published primers amplified multiple dor *crt*.** The expected product was <100 bp. In some but not all cases, it was still possible to determine the sequence for amino acids 72-76 using these reactions.

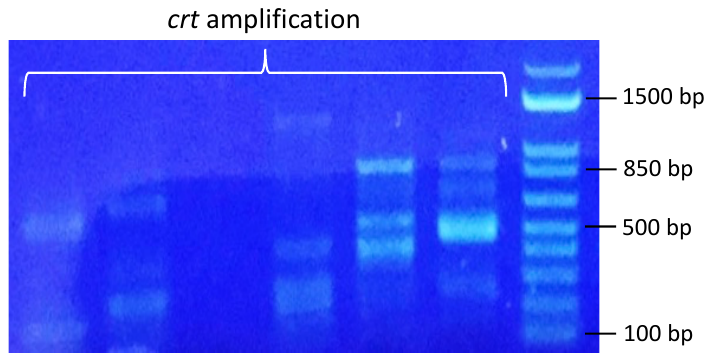
